# Type 2 diabetes mellitus exacerbates vaginal group B *Streptococcus* colonization via impaired mucosal cytokine response

**DOI:** 10.64898/2026.01.08.698441

**Authors:** Clare M. Robertson, Vicki Mercado-Evans, Addison B. Larson, Holly Branthoover, Samantha Ottinger, Marlyd E. Mejia, Zainab A. Hameed, Lindsey A. Gonzalez, Camille Serchejian, Libbie Ogilvie, Jacob J. Zulk, Kathryn A. Patras

## Abstract

Type 2 diabetes mellitus (T2D) is a metabolic disorder that confers increased risk of microbial infections, including those caused by the opportunistic pathogen group B *Streptococcus* (GBS). Asymptomatic GBS carriage in the vaginal tract is a notable reservoir for infection, but the impact of T2D on the vaginal mucosa and GBS colonization is not fully understood. We employed a diet-induced mouse model of T2D paired with vaginal GBS colonization to investigate the impact of diabetes on glucose availability, vaginal microbiome composition, and vaginal cytokine profiles at baseline and in response to GBS. We observed enhanced susceptibility of diabetic mice to GBS vaginal colonization and reproductive tract dissemination. Despite experiencing hyperglycemia, diabetic mice did not exhibit elevated glucose in the reproductive tract. Regarding the vaginal microbiota, diabetic mice had minimal compositional differences with decreased *Mammaliicoccus* being the only significant taxonomic variance. Vaginal cytokine profiling revealed consistently depressed cytokines in diabetic mice, beginning with KC at baseline and expanding to an array of eight pro-inflammatory cytokines post-GBS infection. Pairing cytokine observations with GBS colonization outcomes revealed a correlation between delayed vaginal IL-1α induction and persistent vaginal GBS, suggesting that vaginal cytokine deficiency may contribute to diabetic GBS vaginal colonization. Supplementation with intravaginal rIL-1α was sufficient to resolve GBS burden differences between diabetic mice and non-diabetic controls, confirming that deficient vaginal cytokine responses contribute to diabetic GBS vaginal persistence. These findings advance our understanding of diabetic vaginal mucosal susceptibility to pathogens and support the potential for immunological intervention in the susceptible diabetic population.

**IMPORTANCE:** People with T2D are more susceptible to microbial infections, but there is limited understanding of the mechanisms that drive this vulnerability. One possibility is that T2D enhances colonization of opportunistic pathogens, like GBS, in mucosal reservoirs as a precursor to infection. In this study, we used a diabetic mouse model to test whether diabetes alters the vaginal mucosa to promote GBS colonization. We found that increased vaginal GBS colonization in diabetic mice was not linked to tissue glucose availability or changes to the vaginal microbiome, but instead was associated with impaired vaginal immune responses. These findings provide a foundation for translational approaches to reduce GBS persistence and dissemination in at-risk individuals.

## INTRODUCTION

Group B *Streptococcus* (GBS, *Streptococcus agalactiae*) is a Gram-positive bacterium that asymptomatically colonizes the gastrointestinal and urogenital tracts of approximately 1 in 5 non-pregnant adults(1). While healthy, non-pregnant adults are generally impervious to invasive infection; those with metabolic disorders are more susceptible to a range of manifestations including GBS bacteremia and skin and soft tissue infections(2, 3). In two U.S.-based bacterial surveillance studies, diabetes mellitus and/or obesity were the most common underlying conditions found in GBS non-pregnant adult invasive infections(4, 5). Diabetes mellitus is a metabolic disorder that can be classified into three main types: type 1 (destruction of beta cells, insulin dependent), type 2 (imbalance between insulin production and sensitivity), and gestational (manifests in pregnancy), with type 2 (T2D) being the most common (90% of diabetic cases), impacting more than half a billion people worldwide(6-8). In pregnancy, gestational diabetes elevates risk of rectovaginal GBS colonization by 16% whereas pregestational diabetes (either type 1 or type 2) elevates the risk by 76%(9). Although mucosal GBS colonization is a well-recognized risk factor for invasive disease in the neonatal period(10, 11), the unique diabetic susceptibility to GBS colonization outside of pregnancy, especially in terms of the vaginal reservoir, is underexplored.

Multiple characteristics of T2D could explain susceptibility to GBS vaginal colonization. Three major contributors to susceptibility to pathogens during diabetes are thought to be defective glucose homeostasis, altered composition of the resident microbiota, and aberrant immune recognition and response to pathogens(12-15). Hyperglycemia is thought to increase glucose availability at the mucosal sites where the microbiota resides, which could improve pathogen fitness through direct consumption or modulation of virulence factor expression. Clinically, poor glycemic control is positively associated with GBS colonization risk in women with pregestational diabetes(16); however, a pubertal-onset obesity murine model found no correlations between GBS vaginal burden and systemic glucose intolerance(17). *In vitro* studies have demonstrated wide-spread transcriptional changes in GBS in response to elevated glucose including genes related to virulence and host cell adherence(18, 19). Studies that measure vaginal glucose during diabetes or hyperglycemia are sparse and present conflicting results(20-22), thus it remains unclear whether T2D alters vaginal glucose availability, and in turn, GBS colonization fitness.

The vaginal microbiota is a community of microorganisms that, in a healthy state, exerts control over pathogen invasion and survival via direct competition or indirect modulation of host responses(23). GBS colonization correlates with the absence of beneficial *Lactobacillus* spp. and presence of non-optimal vaginal taxa in both non-pregnant(24, 25) and pregnant(26-29) populations, and many of these synergistic and antagonistic relationships have been established experimentally(30-32). Although taxonomically distinct from the human vaginal microbiota, murine models have also identified correlations between vaginal community composition and GBS colonization success in non-pregnant models(32-34) and GBS infection outcomes in healthy and gestational diabetic pregnancy(35). However, the impact of metabolic disease on the vaginal microbiota is not well-understood. Animal models of pubertal-onset obesity(17) and gestational diabetes(35) have reported modest alterations to vaginal bacterial composition. In humans, vaginal microbial composition in T2D is an active area of research, with recent studies indicating a shift to a higher risk infection-associated, *Lactobacillus* spp. replete profiles(36-39), but the extent to which these alterations influence pathogen colonization is uncharacterized.

Upon vaginal colonization, GBS induces innate immune cytokine and chemokine production by the human vaginal epithelium *in vitro* (such as IL-8 and IL-1β)(40-42) and mice *in vivo* (such as KC, IL-1β, and IL-17)(35, 41), and subsequent cellular immune responses, particularly neutrophils and γδ T cells, are critical in reducing GBS vaginal burdens(35, 43). Diabetes is widely appreciated to coincide with chronic low-grade systemic inflammation, which may be counterproductive for effective pathogen responses(44, 45). Indeed, neutrophils from T2D patients display reduced GBS phagocytosis in experimental hyperglycemia(46). In the context of urogenital immune responses to GBS, diabetic phenotypes appear context dependent. In response to GBS UTI, type 1 diabetic mice display elevated immune cell recruitment(47) whereas type 2 diabetic mice reduced immune cell recruitment(48). In models of gestational diabetes, experimental hyperglycemia in human placental explants reduced cytokine production in response to GBS infection(49), whereas diabetic mice produced exacerbated KC and G-CSF levels when colonized with GBS(35). Whether vaginal immune responses to GBS are altered in T2D has not been established.

Given this site’s potential to serve as a reservoir for GBS to disseminate to other body sites and cause invasive infection, the present study aims to address the role of metabolic parameters, the vaginal microbiota, and mucosal immunity in mediating increased T2D susceptibility to GBS vaginal colonization. Using a diet-induced T2D model in reproductive-age mice, we performed integrated analyses of metabolic phenotypes such as glucose intolerance and vaginal glucose availability, vaginal microbiome profiling, and vaginal cytokine quantification with GBS colonization outcomes. These findings advance our current understanding of the impact of T2D on the vaginal environment and support the potential for immunological intervention to control GBS vaginal colonization in this susceptible population.

## RESULTS

### High-fat, high-sucrose diet causes aberrant metabolism and exacerbates GBS vaginal colonization and dissemination

To assess the impact of diabetes on susceptibility to GBS, we paired murine models of diet-induced T2D and GBS vaginal colonization(50, 51). Adult C57BL/6J female mice were provided a high-fat high-sucrose (HFHS, 45% kcal from fat) diet or a low-fat no sucrose (control, 10% kcal from fat) diet for 12 weeks prior to vaginal inoculation with GBS strain CNCTC 10/84 (serotype V). Every 4 weeks, body mass, vaginal swabs, and vaginal lavage were collected, and a glucose tolerance test was performed at 12 weeks as indicated in **Fig. 1A**. Compared to controls, HFHS-fed mice displayed significantly greater body mass gain as early as 4 weeks after starting diet (**Fig. 1B**), and fasting blood glucose was significantly elevated by 8 weeks on diet (**Fig. 1C**). HFHS-fed mice had impaired glucose tolerance as indicated by heightened blood glucose following glucose bolus and an increased area under the curve compared to control mice (**Fig. 1D-E**).

**Figure 1:**
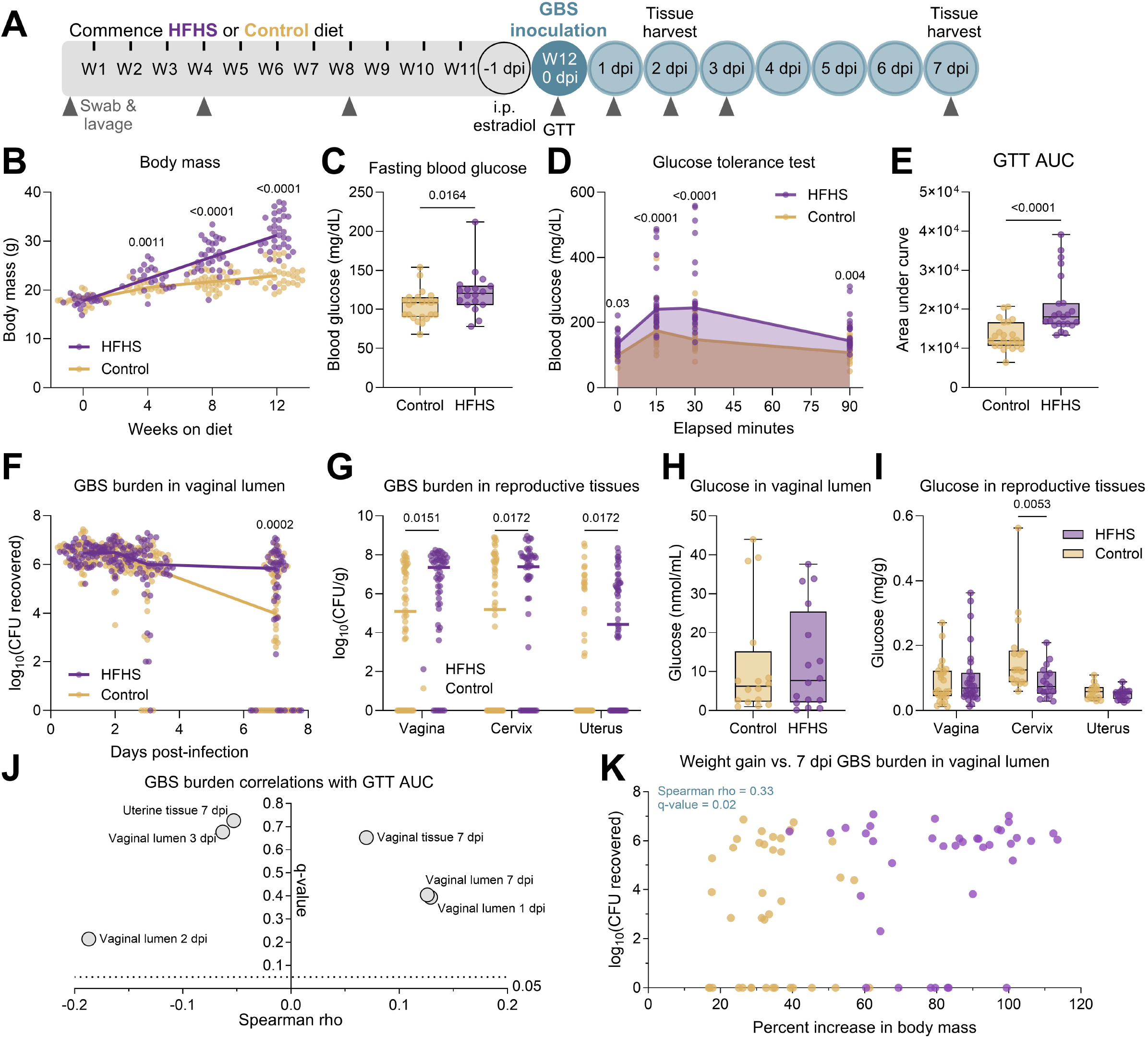
HFHS diet-induced type 2 diabetes exacerbates group B *Streptococcus* vaginal colonization and reproductive tract ascension. **A**) Experimental timeline of type 2 diabetes development on 12 weeks (W) of high-fat high sucrose (HFHS) and control diets and subsequent GBS vaginal colonization and sample collection. **B**) Body mass over time. **C**) Fasting blood glucose at 12 weeks on diet. **D**) Blood glucose concentrations during glucose tolerance test. **E**) Area under the curve (AUC) from glucose tolerance test (GTT). **F**) GBS burdens from post-infection vaginal swabs. **G)** GBS burdens from post-infection homogenized reproductive tissues at 7 days post-infection (dpi). **H)** Glucose concentrations in vaginal lavage fluid after 12 weeks on diet. **I**) Glucose concentrations in post-infection reproductive tissues. **J**) Spearman correlations between GBS swab and tissue burdens and GTT AUC. **K**) Spearman correlation between body mass increase and 7 dpi vaginal swab GBS burdens. *n*=18-34 (B), *n*=18-23(C-E,J), *n*=53-54(F-G), *n*=16(H), *n*=16-28(I), *n*=76(K). Data represent 4 (B-E, I-K), 3 (H), and 8 (F-G) independent experiments. Samples were assayed in duplicate (H,I). Points indicate individual samples, and lines or curves indicate medians. Box and whisker plots show all points and extend from minimum to maximum (C,E,H). Data were analyzed by two-way ANOVA with Benjamini, Krieger and Yekutieli correction and false discovery rate (FDR) set at 5% (B,D, F-G, I), two-tailed Mann−Whitney *U* test (C,E,H), and Spearman correlation with Benjamini-Hochberg FDR correction (J-K).

Compared to control mice, the HFHS-fed group displayed sustained GBS burdens at 7 days post-infection (dpi) in the vaginal lumen, although no differences were seen at earlier timepoints (**Fig. 1F-G**). Furthermore, HFHS-fed mice displayed increased GBS burdens in vaginal, cervical, and uterine tissues at 7 dpi (**Fig. 1H**). To determine the impact of metabolic disease on glucose homeostasis, we quantified glucose in the reproductive tract prior to and following infection. Surprisingly, HFHS-fed mice had no difference in glucose concentration in the vaginal lumen at baseline, nor vaginal or uterine tissue at 7 dpi (**Fig. 1H-I**). In contrast, cervical tissue glucose was significantly decreased in HFHS-fed mice at 7 dpi (**Fig. 1I**). At earlier stages of metabolic disease, there were no differences in colonization outcomes for mice that had only undergone 4 weeks or 8 weeks of diet regimen (**Fig. S1A-D**). There was also no difference in GBS outcomes for mice that underwent a full 12 weeks of diet regimen but only 2 days of GBS colonization (**Fig. S1E-F**). Infections were repeated with a different GBS strain, A909 (serotype Ia) after 12 weeks on diet, but no significant differences in GBS burden between groups were observed in the vaginal lumen or in any of the reproductive tract tissues (**Fig. S1G-H**). Together, these results indicate that the HFHS diet worsens the outcomes of GBS vaginal colonization in a manner dependent on metabolic disease duration, duration of colonization, and GBS strain.

### Elevated body mass gain and post-infection tissue glucose correlate with GBS burden

T2D manifests multiple measurable metabolic features, including impaired glucose tolerance, gain in body mass, and elevated glucose in tissues and fluids(52-54). To clarify the impact of metabolic disease versus direct impact of diet on GBS outcomes, we sought to determine whether any metabolic features correlated with GBS burdens. Glucose tolerance prior to infection did not correlate with GBS burden at any swab timepoint nor in any reproductive tissues (**Fig. 1J**). However, we found that mice that gained more body mass had poor GBS outcomes, indicated by a positive correlation between percent increase in body mass and GBS burdens in 7 dpi swabs and tissues (**Fig. 1K** and **Fig. S2A-B**). To address the relationship between reproductive tract glucose and GBS colonization, we correlated glucose concentrations with GBS burdens. Glucose levels in the vaginal lumen, taken pre-infection, did not correlate with GBS burden at any swab timepoint (**Fig. S2C**), indicating that vaginal glucose availability does not impact infection outcomes. At 7 dpi, glucose levels in vaginal tissue did not significantly correlate with vaginal tissue GBS burdens, but cervical and uterine glucose negatively correlated with GBS burdens at their respective sites (**Fig. S2D**). These data suggest that GBS glucose consumption is augmented in the cervix and uterus compared to vagina, or that glucose limitation benefits GBS spread to the cervix and uterus. The results also reveal no increase in glucose levels in the diabetic female reproductive tract, discordant with elevated levels seen systemically and at other mucosal sites(55, 56).

### Diabetic mice have minimal changes to the vaginal microbiome

The vaginal microbiome is implicated in susceptibility to urogenital infection, but little is known about the vaginal microbiome in the context of T2D(36-39). To assess changes in the microbiome composition at successive stages of metabolic disease development, we profiled the vaginal microbiome of HFHS-fed and control mice at 0, 4, 8, and 12 weeks on the diet. Vaginal communities in both groups were primarily dominated by *Staphylococcus xylosus* at all timepoints, with less frequent dominant taxa including *Staphylococcus equorum, Enterococcus faecalis, Mammaliicoccus lentus, Streptococcus agalactiae* (GBS) and *Escherichia* spp. (**Fig. 2A**). The spontaneous emergence of GBS was not retained in any individual mouse across more than one timepoint, indicating that GBS presence in the vaginal microbiota of these mice was transient (**Table S1**). At all timepoints, no significant differences in alpha diversity metrics, including amplicon sequence variants (ASVs) or Shannon Entropy, were detected (**Fig. 2B-C**). At 4- and 8-week timepoints, no differentially abundant genera or species were detected (**Table S1**). At 12 weeks on the diet, differential abundance analysis via ANCOM-BC2 revealed a decrease in *Mammaliicoccus* genus in the HFHS-fed group (**Fig. 2D**). Two alternative methods, Mann-Whitney and mixed linear modeling, did not identify differentially abundant genera (**Table S1**). Additionally, no species or ASVs were significantly differentially abundant by any method (**Fig. 2E, Table S1**). Despite frequent sequencing- and cultivation-based detection of murine vaginal *Enterococcus* in prior studies using the same vendor and vivarium(32, 35, 57), in this study, detection of vaginal *Enterococcus* via sequencing was rare (**Table S1**), and no differences in vaginal swab or tissue *Enterococcus* detection rate or CFU were observed between diet groups (**Fig. S3A-D**). However, *Enterococcus* burden positively correlated with GBS burden at later timepoints (3 and 7 dpi) in vaginal swabs and in all 3 reproductive tissues at 7 dpi (**Fig. S3E-F**).

**Figure 2:**
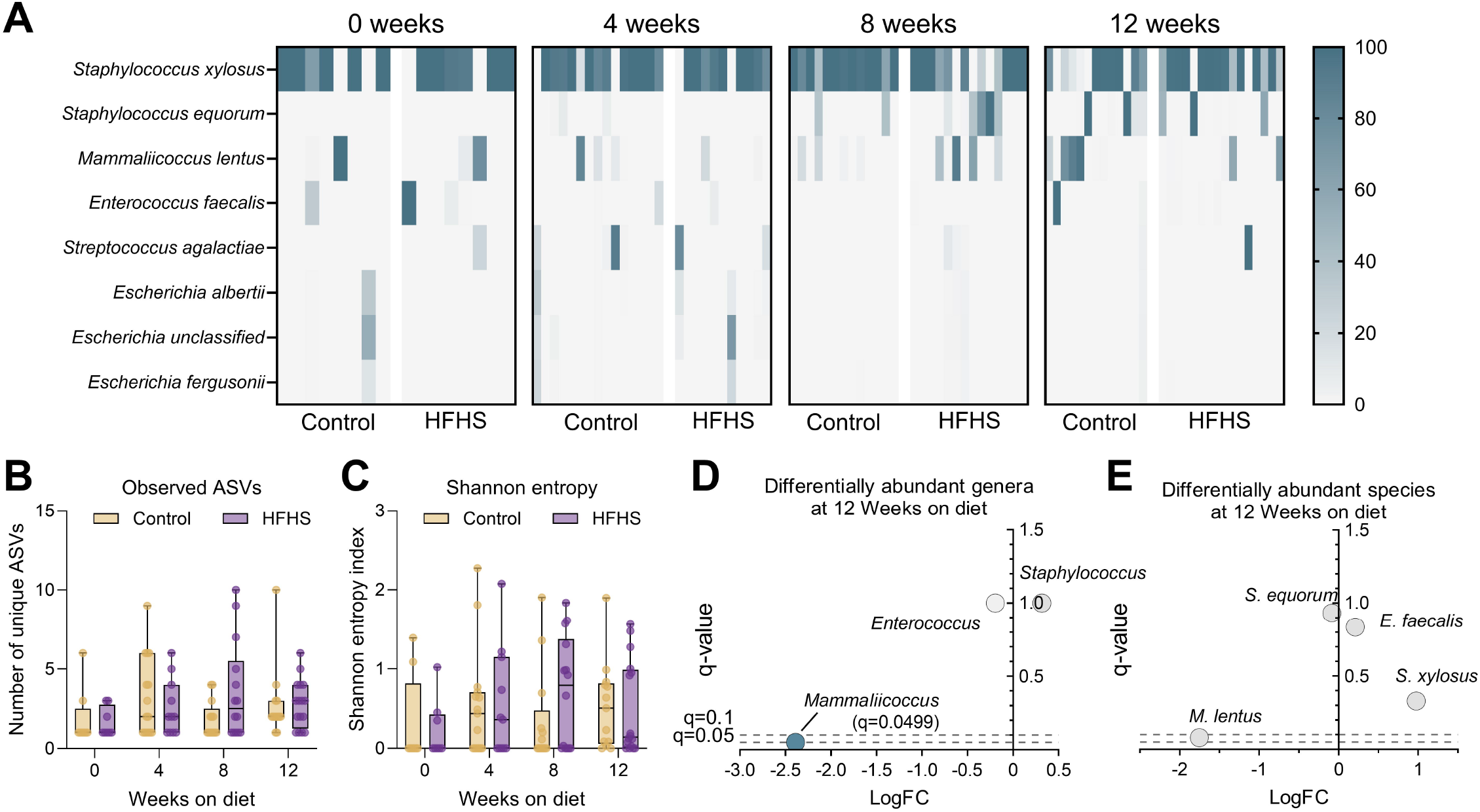
Twelve weeks of HFHS diet does not alter vaginal microbiome bacterial diversity but decreases the relative abundance of *Mammaliicoccus*. **A**) Heatmap of relative abundances for bacterial genera/species in individual mice at 0, 4, 8, or 12 weeks on diet. **B**) Total unique observed amplicon sequence variants (ASVs) in each group at 0, 4, 8, or 12 weeks on diet. **C**) Shannon entropy indexes) in each group at 0, 4, 8, or 12 weeks on diet. **D**) Differentially abundant genera in HFHS diet group compared to control diet after 12 weeks. **E**) Differentially abundant species in HFHS diet group compared to control diet after 12 weeks. *n*=8-16 (A-C), *n*=13-16(D-E). Data represent 2 independent experiments. Heatmap columns represent individual mice (A). Box and whisker plots show all points, representing individual samples, lines indicate medians, and whiskers extend from minimum to maximum (B-C). Data were analyzed two-way ANOVA with Benjamini, Krieger and Yekutieli correction and false discovery rate (FDR) set at 5% (B-C) or ANCOM-BC2 with diet as a fixed effect (D-E).

### Diabetic mice have aberrant vaginal cytokine responses to GBS

Another potential mechanism by which type 2 diabetes may confer increased susceptibility to GBS is through alterations to reproductive tract immune responses. To test this possibility, we quantified 23 cytokines in vaginal lavages collected at 4, 8, and 12 weeks on diet as well as 2 dpi and 7 dpi, timepoints with established innate immune induction in this model(40, 41). Cytokines that were detected in at least 2 samples per group were retained for analysis for each timepoint (**Fig. S4, Table S2**). While no significant differences were seen at 4 or 8 weeks on diet, by week 12, HFHS-fed mice displayed lower levels of chemokine KC (**Fig. 3A-B, Fig. S4**). At 2 dpi, eight cytokines were significantly lower in the HFHS-fed group compared to the control group, including MCP-1 (**Fig. 3A, C**). No significant cytokine differences were observed at 7 dpi even though GBS burdens were higher in HFHS-fed mice at that timepoint, but we did note differential IL-1α fluctuation between HFHS and control mice over the course of infection (**Fig. 3D**). We then correlated cytokine levels with metabolic markers of glucose intolerance, body mass, and weight gain. Glucose intolerance negatively correlated with pre-infection KC levels at 12 weeks on diet and with post-infection MCP-1 levels at 2dpi (**Fig. 3E-F**). Additionally, we observed a positive correlation between 4-week MCP-1 and 12-week GTT AUC and inverse correlation between 4-week IL-4 and 12-week GTT AUC, but these findings did not achieve statistical significance (**Table S2**). Taken together, these results indicate that the HFHS diet impairs local vaginal immune signaling in response to GBS challenge and suggests broad impacts on pathways relating to both innate and adaptive immunity.

**Figure 3:**
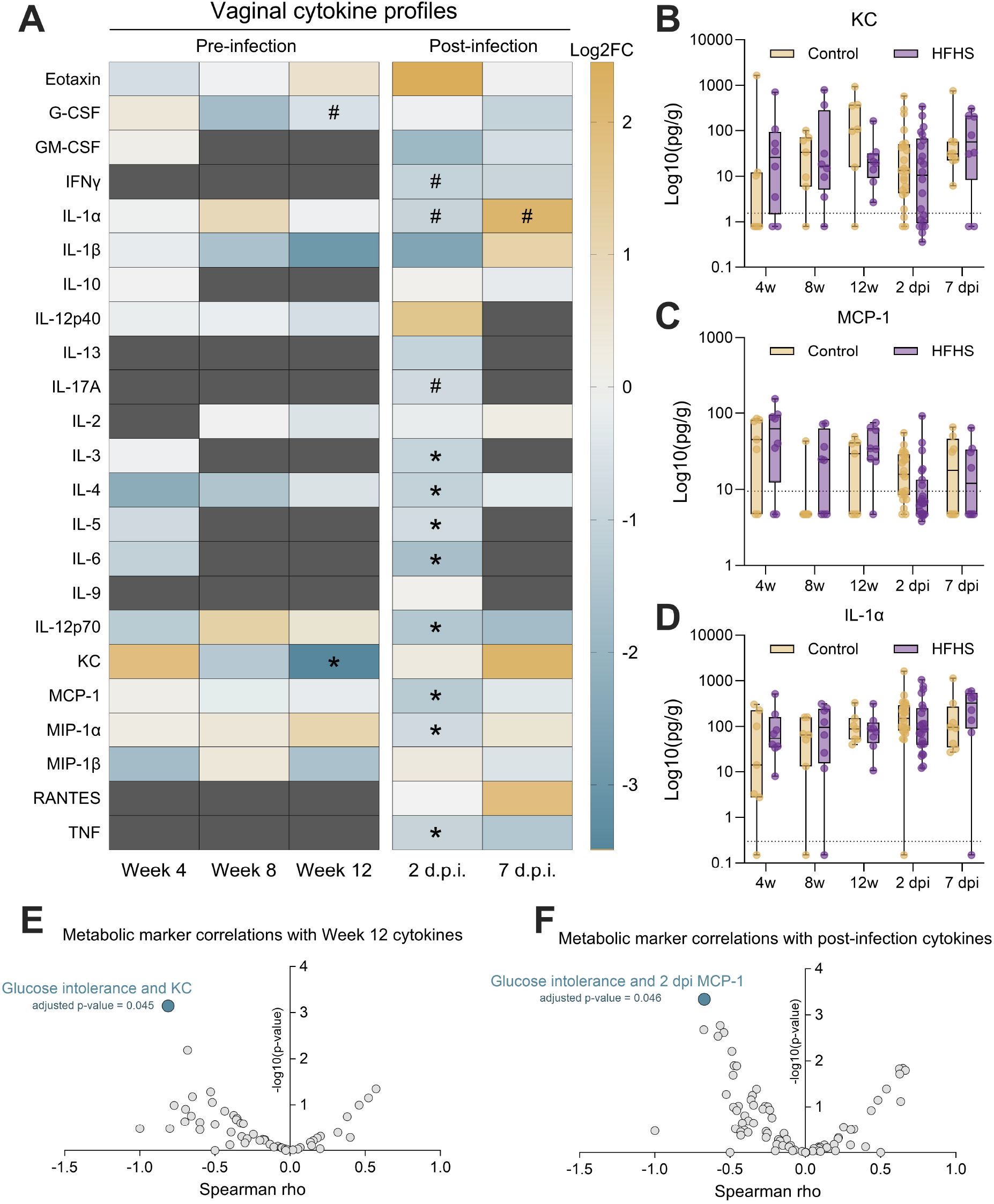
HFHS diet suppresses a baseline vaginal chemokine and broadly suppresses early vaginal cytokine response to GBS. **A**) Vaginal lavage fluid cytokine profiles pre-and post-infection, displayed as log2FC of HFHS group median over control group median. Dark grey indicates timepoints where at least one group had less than 2 samples meeting the lower detection limit of that cytokine. Vaginal lavage concentrations of KC (**B**), MCP-1 (**C**), and IL-1α (**D**) normalized to total protein. Spearman correlation results between metabolic markers and baseline (**E**) or post-infection (**F**) vaginal cytokine concentrations. *n*=7-16 (A-D). Data represent 1 (pre-infection and 7 dpi timepoints) or 3 (2 dpi timepoint) independent experiments. Box and whisker plots show all points, representing individual samples, lines indicate medians, and whiskers extend from minimum to maximum, with dotted lines indicating the lower limit of detection (B-D). Data were analyzed by two-tailed Mann−Whitney *U* test (A-D) or Spearman correlation with Benjamini-Hochberg FDR correction (E-F). ^#^*p*<0.1, \**p*<0.05.

### Exogenous IL-1α treatment ablates diabetic susceptibility to GBS

We next sought to confirm whether immune deficiencies directly contribute to the increased susceptibility to GBS seen in diabetic mice. To clarify which cytokines were linked to infection outcomes, we performed Spearman correlations between GBS burden and cytokine concentrations at paired timepoints. IL-1α at 7 dpi was the only significant cytokine that correlated with GBS CFU (**Fig. 4A-B, Table S2**). Because of this connection between IL-1α and GBS burden, and general pattern of reduced cytokines in HFHS-fed mice, we selected this broadly acting cytokine for an intervention study. We supplemented mice with topical vaginal recombinant IL-1α beginning one day prior to GBS infection and daily throughout the infection time course (**Fig. 4C**). Treatment with IL-1α abrogated differences in GBS burdens in the vaginal lumen and reproductive tract tissues at 7 dpi and rates of GBS uterine ascension between diabetic and control groups (**Fig. 4D-F**). Taken together, these results indicate that increased GBS colonization and dissemination in diabetic mice can been absolved by supplementing with a broadly acting cytokine at the site of colonization.

**Figure 4:**
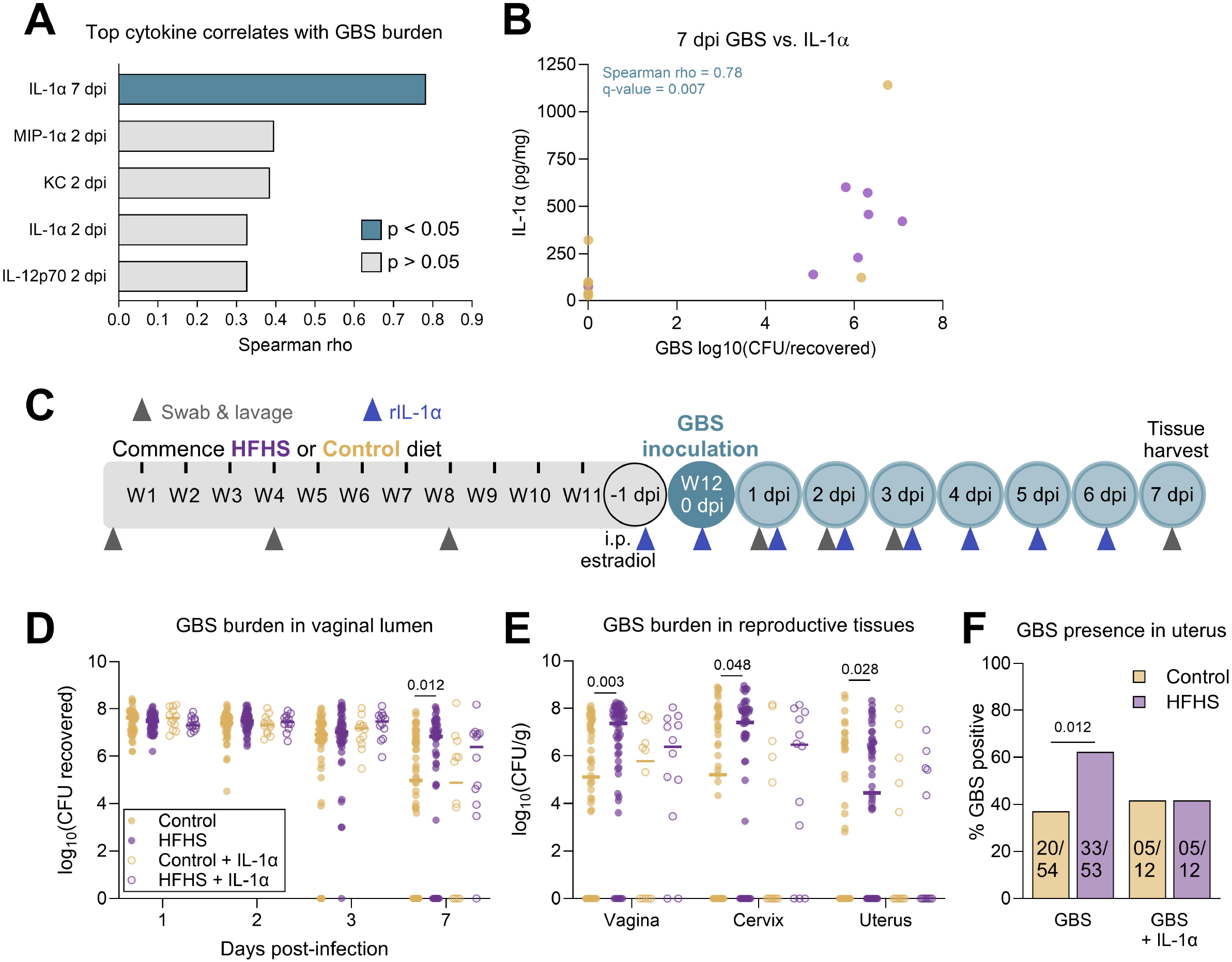
IL-1α supplementation resolves elevated endpoint GBS burdens and uterine ascension in HFHS-fed mice. **A**) Spearman correlations between vaginal cytokine concentrations and vaginal GBS burdens at paired timepoints. The cytokines with the 5 strongest rho values are displayed. **B**) 7 dpi vaginal swab GBS burdens plotted with 7 dpi vaginal IL-1α. **C**) Experimental timeline of type 2 diabetes development on HFHS diet and subsequent rIL-1α supplementation paired with GBS vaginal colonization. **D**) GBS burdens from post-infection vaginal swabs. **E**) GBS burdens from post-infection reproductive tissues. **F**) Proportions of mice in each group with GBS detected in uterine tissue. *n*=7-8 (A-B), *n*=12-54 (D-F). Data in D-E represent data from Figure 1F-G shown for comparison with rIL-1α treatment groups. Data represent 1 (A-B), 2 (D-F, groups receiving rIL-1α), or 8 (D-F, groups not receiving rIL-1α, two of which were performed in tandem with rIL-1α treatment) independent experiments. Data were analyzed by Spearman correlation with Benjamini-Hochberg FDR correction (A-B), two-tailed Mann−Whitney *U* test with Benjamini, Krieger and Yekutieli correction and FDR set at 5% (D-E), or two-sided Fisher’s exact test (F).

## DISCUSSION

Although type 2 diabetes is a leading risk factor for GBS invasive disease in non-pregnant adults(5), biological drivers of this increased susceptibility are not fully known. In this study, we found that a diet-induced murine model of T2D exhibits increased GBS vaginal persistence and ascension to the upper reproductive tract, The vaginal microbiome was minimally altered in diabetic mice, suggesting it has negligible impact on GBS outcomes in this model. Conversely, we found that HFHS-fed mice failed to mount a vaginal pro-inflammatory response to GBS, and that certain cytokine responses appear delayed. We further showed that stimulating an inflammatory response early during colonization prevents the heightened susceptibility to GBS in HFHS-fed mice. Together, these findings reveal impaired mucosal immune responses and enhanced pathogen colonization as potential precursors of GBS invasive disease in diabetic individuals.

We selected a diet-induced model as a relevant simulation of T2D, considering the majority of T2D cases are attributable to diet(58), and there is an increasing global prevalence of highly processed diets high in fats and sugars(59). Using a 7-week course refined diet model, Megli et al. recently described increased GBS persistence in mice fed any refined diet - either high fat, low soluble fiber or low fat, low soluble fiber - compared to regular chow(17). Similar to Megli et al, we found no correlations between glucose intolerance and GBS burdens, however, divergently, we found that body weight gain significantly correlated with GBS vaginal levels at day 7. There are several experimental differences that may explain this discordance. First, the length of dietary exposure (7-week vs. 12-week) may accentuate phenotypes in our study. Additionally, Megli et al supplied experimental diets over puberty, whereas our study exclusively treated mice post-puberty, which may have altered the extent of metabolic disease. Finally, dietary formulations, although similar in terms of fiber (5-6%), differed in terms of high fat (60% vs. 45% energy from fat) and sucrose content (9% vs. 21%) between their study and this study, respectively. Collectively, these findings suggest that both dietary components and metabolic metrics impact GBS susceptibility, aligned with clinical observations of obesity and sugary drink intake as risk factors for GBS vaginal colonization(60-62).

Elevated hemoglobin A1c (HbA1c), an indicator of sustained hyperglycemia, is associated with increased risk for vaginitis and genital infections including HPV and Candidiasis(63-65), as well as colonization by pathogens including GBS and other pathobionts(16, 66, 67). Although diabetic patients display elevated glucose in mucosal secretions such as saliva and tears(55, 68, 69), few citations exist wherein diabetic vaginal glucose is empirically measured(20, 21). One human study quantified vaginal glucose in primarily non-diabetic patients before and during glucose tolerance testing and discovered a surprising decrease in vaginal glucose after the oral glucose bolus, independent of HbA1c, plasma glucose, or BMI(20). These findings suggest that, during a hyperglycemic state, excess glucose in the vaginal mucosa is effectively metabolized or stored, reducing its detection in the lumen. Further studies are needed to determine the impact of hyperglycemia on the fate of glucose in the reproductive tract. Excess glucose may accumulate inside differentiated keratinocytes(70), a phenomenon which could select for pathogens with capacity for cytotoxicity or intracellular invasion. Alternatively, it may be metabolized by microbiota directly or after conversion to glycogen(71), or non-enzymatically incorporated into advanced glycation end products, with myriad downstream impacts on tissue physiology and architecture(72, 73).

In our model, vaginal lavage and tissue glucose levels were similar in diabetic and control mice. Moreover, cervical tissue glucose levels were decreased in HFHS-fed mice. This may be attributable to GBS utilization of glucose, as GBS burdens negatively correlated with glucose concentrations in cervix. However, the same negative correlation between GBS and glucose was observed in uterine tissue, despite glucose levels being similar in uterine tissue of diabetic and control mice. Compared to vaginal and cervical glucose levels, uterine glucose levels were lower overall. These disparate findings in the lower versus upper reproductive tract suggest that GBS may alter its metabolism as it adapts to tissue niches. In line with this, multiple GBS carbon utilization genes are differentially expressed between vaginal and uterine tissues in gestational diabetic mice compared to controls including a putative GBS glycosyltransferase *yfhO* critical for GBS uterine ascension(35). In contrast to our findings, increased vaginal glucose was reported in a type 1 diabetic mouse model(22), suggesting alternative physiological impacts on tissue glucose homeostasis across diabetic manifestations.

The human vaginal microbiome has been profiled in gestational diabetes mellitus(74-77), and more recently in T2D(36-39). Although there is heterogeneity in specific taxonomic findings, collectively these studies suggest diabetes-related reduction in health-associated *Lactobacillus* spp. and enrichments in infection-associated bacterial taxa. Some of these taxa, such as *Enterococcus* and *Escherichia*, co-occur with GBS *in vivo* (35, 78-81) and have the capacity to behave synergistically with GBS *in vitro* (82). Their enrichment may partially explain the increased susceptibility to GBS colonization in diabetics(9, 16). A key limitation of murine models is the inability to replicate the vaginal community composition of humans, even though overall low alpha diversity and dominance by one or two taxa are shared between species(32, 33, 57). Megli et al. used a conventional mouse model to investigate microbiome dynamics before and during long-term GBS vaginal colonization, wherein they found that mice on a high fat diet had increased vaginal *Streptococcus* and *Enterococcus* and decreased *Staphylococcus* at baseline and at multiple post-infection timepoints(17). HFHS mice in our current study had decreased relative abundance of *Mammaliicoccus*, a close relative to *Staphylococcus* in the family *Staphylococcaceae*, suggesting shared constraints on a similar microbial niche. However, *Streptococcus* and *Enterococcus* were not detected in many of our mice. Demirel et al. profiled vaginal microbiomes of hyperglycemic rats and found no significant changes(83). These dissimilar findings reflect that inherent differences in species, vendors, and/or animal facilities play a role in bacterial compositions and thus influence microbiome findings downstream.

T2D is often considered a chronic inflammatory condition, with sustained elevation of immune markers such as C-reactive protein and IL-6(44, 84). Other studies have shown the predictive power of pro-inflammatory and immunoregulatory cytokine levels for a future T2D diagnosis(44, 85-88). While systemic baseline inflammation is well-documented, T2D impacts on mucosal immune responses, particularly in the context of infection, are more sparsely described. When we profiled vaginal cytokines in our HFHS mouse model, we discovered a broad suppression in many pro-inflammatory cytokines compared to control mice. These effects appeared as early as 4 weeks on diet and were most prominent during GBS challenge. Our results may be sex-specific or specific to certain epithelial sites. Of the few studies that investigated sex differences in systemic diabetic inflammation, in adolescent humans with T1D and a rat model of T2D, only males exhibit increases in inflammatory cytokines, whereas females exhibit decreases in these same inflammatory cytokines(89, 90). Further, a study on female mini-pigs showed that long-term high fat diet lowered serum IL-1β, IL-10, and IL-4, suggesting a general decrease in both pro- and anti-inflammatory cytokine-mediated immune signaling(91). Regarding site specificity, diabetic mouse models of GBS infection generate distinctive inflammation responses depending on the infected tissue. In a T2D urinary tract infection (UTI) model, both male and female db/db mice had suppressed pro-inflammatory cytokines and reduced immune cell infiltration in the bladder(48). In contrast, models of diabetic lung, wound, arthritis, T1D UTI, and systemic infection show that diabetic mice have increased pro-inflammatory cytokines and immune cell infiltration or upregulated immune cell infiltration pathways(47, 92-94). Finally, it is possible that multiple host metabolic factors shape the mucosal immune landscape in diabetes, leading to this collection of diverging results among distinct diabetic disease types and host states. Indeed, gestational diabetic mice of the same genetic background and fed the same diet as this current study displayed several increased inflammatory vaginal cytokines compared to non-diabetic pregnant controls upon GBS challenge(35).

Our observed cytokine phenotypes indicate that immune impairment develops over the course of metabolic disease progression in the T2D vaginal tract. KC and MCP-1 are implicated in the development of obesity and insulin resistance(95, 96), and we observed a positive, but non-significant, correlation between 4-week MCP-1 and subsequent glucose intolerance. At later timepoints, we observed inverse correlations between vaginal levels of these chemokines and glucose intolerance. It is also possible that diabetes alters immune response kinetics. GBS stimulates robust IL-1α induction at the uroepithelium(97, 98), gravid reproductive tract(35), and choriodecidua(99), and here, we found that GBS burdens were highly correlated with vaginal IL-1α at 7 dpi, primarily driven by diabetic mice. When we supplemented with exogenous rIL-1α, heightened susceptibility to GBS was ablated in HFHS-fed mice, suggesting that immune stimulation is sufficient to improve vaginal GBS colonization outcomes in diabetic mice. Glucose metabolism and IL-1α production have been linked in other tissues; IL-1α increases glucose uptake in muscle(100) and elevated glucose stimulates IL-1α production in renal epithelial cells(101). In invasive infection with closely related group A *Streptococcus*, IL-1α coordinated liver metabolic adaptation and tolerance to infection(102); however, the impact of IL-1α, whether endogenous or exogenous, on reproductive tract tissue glucose homeostasis and downstream effects on immunity remain uncharacterized. Longitudinal profiling of systemic and local metabolism and immunity and immune cell behavioral assays (e.g., chemotaxis/killing) are needed to confirm these observations.

It is necessary to note the limitations of our study. Firstly, diet-induced models of metabolic disease cannot untie the impacts of dietary components from the impacts of metabolic disease. With its long-term 12-week diet time course, our study design minimizes the impact of early host and microbiota adaptation to dietary components. Our study would be complemented by replication studies using genetically predisposed (e.g., db/db) or chemically induced (e.g., STZ) models. Second, our experiments were conducted primarily with one GBS strain (CNCTC 10/84) which demonstrated increased colonization *in vivo*, whereas another strain, A909, did not exhibit differential colonization in diabetic mice. Thus, our findings may only be applicable to certain GBS genetic backgrounds. CNCTC 10/84 has a mutation in the *covR* promoter(103), the regulatory component of the covRS system that coordinates GBS virulence and response to glucose(18). *De novo covRS* mutations are observed in a mouse diabetic wound model(93) and in human vaginal isolates(104), supporting the relevance for modeling host-GBS interactions in the diabetic reproductive tract. Third, our microbiome profiling experiments are constrained due to the limited suitability of the murine vaginal microbiome as a model of the human vaginal microbiome. Whereas the majority of human vaginal microbiomes are dominated by a *Lactobacillus* species, conventional mice are very rarely dominated by any *Lactobacillus* or closely related genera(32, 33). The potential to determine vaginal microbiome phenotypes of type 2 diabetes will likely be greatest in human studies and in humanized models of the vaginal microbiome(32, 105). Lastly, we have identified a potential sex difference or tissue specificity in diabetic immune phenotypes, but our all-female study design and focus on the female reproductive tract inherently precludes traditional sex-based comparisons. Future studies should also investigate mechanisms of sex differences in the diabetic immune response, including sex hormones and genetic components.

In summary, we showed that dysregulated immune responses to group B *Streptococcus* at the vaginal mucosa contribute to increased bacterial burdens during colonization. Our findings provide an explanation for increased GBS vaginal colonization in type 2 diabetics and highlight that deficient immunity may be an important but underexplored phenotype in females and/or at mucosal sites. Future investigation should focus on pharmaceutical immune modulation approaches to control GBS colonization in the vulnerable diabetic host. Furthermore, because type 2 diabetics are susceptible to an array of urogenital pathogens, the impact of diabetes-associated immune suppression should be investigated with regard to other taxa, such as *Klebsiella, Escherichia coli*, and *Candida* spp. In this population, immunostimulatory therapies may be a viable alternative to antibiotic-based treatments in the age of increasing recurrent infection and antibiotic resistance.

## MATERIALS AND METHODS

### Animals

Female wild-type C57BL/6J mice from Jackson Laboratories (strain code 000664) were used for all experiments. Mice were housed at a maximum of 4 animals per cage and given food and water ad libitum. A 12-hour light cycle was used. Mice were purchased at 6 weeks old and randomly assigned to a diet group. The two groups were fed a high-fat high-sucrose (HFHS) diet (D12451, Research Diets Inc., 45% kcal fat, 17% kcal sucrose) or a control low-fat no sucrose diet (control) (D12450K, Research Diets Inc., 10% kcal fat, <1% kcal sucrose) for up to 12 weeks prior to GBS colonization, until the experimental endpoint, up to a total of 13 weeks. A subset of mice were originally used for timed pregnancy experiments and were cohabitated with males for 3 days after 1 week on diet. Only mice that did not become pregnant were used for the present study. All animal protocols and procedures were approved by the BCM Institutional Animal Care and Use Committee.

### Assessment of metabolic markers

To determine body mass, mice were weighed on a scale at baseline prior to diet, then at 4 weeks, 8 weeks, and 12 weeks after starting diet. To determine fasting blood glucose, food and water were withheld for 4 hours, then blood glucose was measured using a glucometer (ReliOn Prime Blood Glucose Monitoring System) via tail venipuncture. To determine glucose tolerance, food and water were withheld for 4 hours, then baseline fasting blood glucose was measured at the 0-minute timepoint. Mice were then given an intraperitoneal injection of 1mg glucose in PBS per 1g body mass in a 100µL volume. Blood glucose was then measured at 15, 30, and 90 minutes post glucose bolus.

### GBS vaginal colonization

Mice were colonized as described previously at 4, 8, or 12 weeks after beginning the diet(51). One day prior to GBS infection, mice were injected intraperitoneally with 0.5mg 17β-estradiol (Sigma-Aldrich) suspended in 100µL sesame oil. *Streptococcus agalactiae*, or group B *Streptococcus* (GBS) was grown in Todd-Hewitt broth (THB) (Hardy Diagnostics, Santa Maria, CA) at 37□°C in aerobic, static conditions. Wild-type clinical GBS isolates CNCTC 10/84 (ATCC 49447, serotype V) or A909 (ATCC BAA-1138, serotype Ia) were grown overnight in 3mL THB, then sub-cultured by adding 0.3mL overnight culture to 2.7mL fresh THB and grown to log phase at an OD600 of 0.4-0.6. GBS was pelleted then resuspended in PBS. Each mouse was inoculated vaginally with 10^7^ CFU of GBS suspended in 10 µL using a gel-loading pipette tips as described previously(51).

### GBS quantification in vaginal lumen

Vaginal swabs were collected as described previously(51). Briefly, sterile nasopharyngeal swabs (Puritan) were placed in 100µL sterile PBS to wet the swab tip, then inserted in the vaginal opening. Swabs were rotated 4 times clockwise and 4 times counterclockwise, then removed and placed back in sterile tubes of PBS. Blank samples were generated during each sample collection by exposing a sterile swab to the laminar air flow hood environment for several seconds, then re-submerging the swab in the tube of PBS and processing with identical methods as other swab samples. Swabs were secured inside tubes and tubes were vortexed for 30 seconds to dislodge bacteria from swab tip. Samples (10µL) were used for serial dilution and plating on selective chromogenic agar media (CHROMAgar StrepB, DRG International, Inc.). Purple/mauve colonies were counted to quantify GBS CFU and dark blue colonies were counted to quantify *Enterococcus* CFU.

### GBS quantification in reproductive tissues

At the experimental endpoint, mice were sacrificed, and the vagina, cervix, and uterus were harvested from each mouse. Whole tissues were placed in pre-weighed 2mL screwcap microcentrifuge tubes containing 500µL PBS and 1.0 zirconia-silicate beads. Tubes were weighed again after addition of tissues and stored in ice until placed in a Roche Magnalyser bead beater. Tissues were homogenized at 6000 rpm for 60 seconds. 10µL of each homogenized tissue sample was used for serial dilution and plating on CHROMAgar StrepB. Purple colonies were counted to quantify GBS CFU and dark blue colonies were counted to quantify *Enterococcus* CFU. Cardiac blood was harvested from each mouse and plated un-diluted on CHROMAgar StrepB to confirm the absence of GBS bacteremia.

### Vaginal microbiota 16S rRNA sequencing

Vaginal lumen swabs were collected as described above, and samples were stored at -20°C until microbial DNA was extracted using Quick-DNA Fungal/Bacterial Microprep Kit (Zymo Research). DNA was eluted in 20μL molecular biology grade water. At SeqCoast Genomics (Portsmouth, NH, USA), a sequencing library was generated by amplifying the v3/v4 region of the 16S rRNA gene via PCR using the primer pair 341F and 806R. using the Zymo Quick-16S Plus NGS Library Prep Kit. The DNA library was sequenced on an Illumina NextSeq2000 platform using a 600-cycle flow cell kit to produce 2x300bp paired reads. 30-40% PhiX control (unindexed) was spiked into the library pool to support optimal base calling of low diversity libraries on patterned flow cells. Read demultiplexing, read trimming, and run analytics were performed using DRAGEN v4.2.7 software on the NextSeq2000. Reads were combined into a single FASTA file, and the rest of the sequencing analysis was performed in QIIME2 v2024.2 unless otherwise noted. Sequences were joined and trimmed to 300 bp, then denoised using the DADA2 plugin. OTUs were assigned taxonomy by mapping reads to the Greengenes2 database v 2022.10 with a similarity cutoff of 99%. Decontamination was performed by filtering out ASVs that appear in blanks or in <5% of samples and removing known contaminants from the DNA extraction process (**Table S1**). Decontaminated sequences were returned to QIIME2 for community diversity analyses.

### Quantification of cytokine levels in vaginal lumen

Pre-infection vaginal lavages were collected at 4 weeks, 8 weeks, or 12 weeks on diet immediately before GBS infection (0 dpi). Post-infection vaginal lavages were collected at 2 dpi or 7 dpi. Lavages were performed by pipetting 10µL sterile PBS into the vagina of each mouse, mixing by pipet 4 times, then diluting the sample 5-fold in sterile PBS. Lavage samples were subjected to a 23-plex ELISA assay (Bio-Rad, Cat. No. M60009RDP) to quantify IL-1α, IL-1β, IL-2, IL-3, IL-4, IL-5, IL-6, IL-9, IL-10, IL-12 (p40), IL-12 (p70), IL-13, IL-17A, Eotaxin, G-CSF, GM-CSF, IFN-γ, KC, MCP-1, MIP-1α, MIP-1β, RANTES, and TNF-α. Samples were thawed on ice, then centrifuged at 10,000*g* for 10 minutes. Supernatant was diluted 1:10 in Bio-Rad sample diluent, then processed according to manufacturer protocol. Fluorescence was detected on a Luminex MAGPIX instrument and data was generated with Luminex xPONENT software for Magpix, version 4.2 build 1324. Data was analyzed with Milliplex Analyst, version 5.1.0.0 standard, build 10/27/2012. Log curves were fit to the mean fluorescent intensity data to calculate cytokine concentrations, with bovine serum albumin (BSA) standards as a reference. Cytokine data was normalized to total protein levels measured from each sample using a BCA assay (Pierce).

### Exogenous IL-1α intervention experiments

Mice were vaginally inoculated with 80 pg of recombinant mouse IL-1α (BioLegend, Cat. No. 575002) resuspended in 10µL sterile PBS. Mice were treated every day beginning 1 day prior to infection, with the last treatment the day prior (6 dpi) to the endpoint. Mock intervention treatment was vaginal inoculation with 10µL sterile PBS. All mice were then subjected to GBS vaginal colonization and GBS burden monitoring as described above.

### Statistical analyses

All experiments were completed in at least duplicate unless otherwise indicated with results combined prior to analyses. Experimental sample size (n) and exact tests used are indicated in figure legends. Data normality was tested using the D’Agostino-Pearson normality test. Data were analyzed according to the caption under each corresponding figure. For all experiments, a p-value or q-value of <0.05 was considered statistically significant. Statistical analyses were performed using GraphPad Prism v9.5.1, R v4.3.3, or python v3.9.2.

## Data and code availability

Raw sequencing files can be found in the NCBI Sequence Read Archive (SRA) and associated BioProject, accession PRJNA1394477. Code for data analysis can be found in the Patras Lab Github: https://github.com/PatrasLab/T2D_GBS_mouse_vaginal_colonization_manuscript.

## ACKNOWLEDGEMENTS

We are grateful to the vivarium staff at BCM for animal husbandry. Multiplex cytokine analysis was performed by the BCM Antibody-based Proteomics Core, supported in part by CPRIT Proteomics & Metabolomics Core Facility Support Award (RP210227) and NCI Cancer Center Support Grant (P30CA125123). We thank Shixia Huang, Zhongcheng Shi, and Yuan Yao for their excellent technical assistant in performing experiments, data preliminary analyses and QC, and project consultation. Data analysis for 16S sequencing and all correlation analyses were performed on the HPC cluster that is managed by the Biostatistics and Informatics Shared Resource (BISR) and supported by an NCI P30-CA125123 and Institutional funds from the Dan L Duncan Comprehensive Cancer Center and Baylor College of Medicine. This research was supported by NIH R01 (DK128053), NIH R21 (AI173448), and Burroughs Wellcome Next Gen Pregnancy Initiative (NGP10103) grants to KAP. CMR, VME, HB, SO, MEM, JJZ, and HB were supported by NIH F31 training grants (DK138748, AI167547, HD117458, HD111236, AI167538, and DK136201, respectively). HB was also supported by an NIH T32 training grant for a portion of the project (AI055449). LO was supported by a NIH R25 Post-baccalaureate Research Education Program (PREP) award for a portion of the project (GM069234).

## AUTHOR CONTRIBUTIONS

Conceptualization: CMR, KAP; Methodology: CMR, VME, HB, SO, MEM, KAP; Investigation: CMR, VME, ABL, HB, SO, MEM, ZAH, LAG, CS, LO, JJZ, KAP; Formal analyses: CMR; Funding acquisition: KAP; Writing – original draft: CR, KAP; Writing – review and editing: all authors.

