## Supplemental Material for "Type 2 diabetes mellitus exacerbates vaginal group B *Streptococcus* colonization via impaired mucosal cytokine response"

**Contents:**

Supplemental Figures 1-4

Captions for Supplemental Tables 1-2

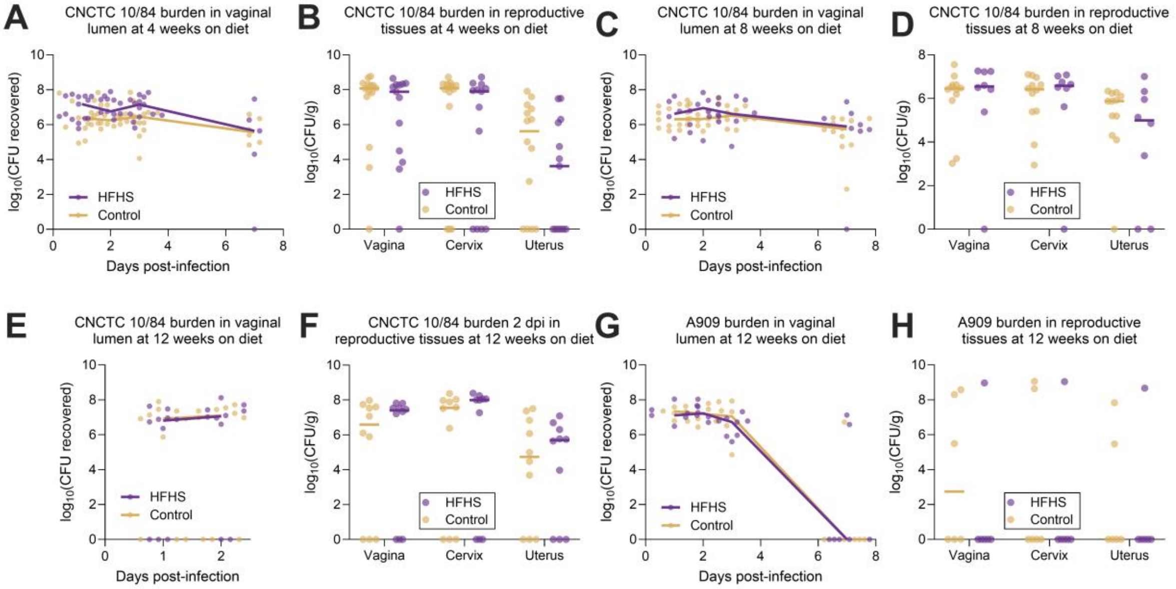


**Figure S1: GBS burdens in mice with shorter diet regimens and alternative GBS strains**. Mice were subjected to a 4-, 8-, or 12-week diet course and GBS burdens were quantified from post-infection vaginal swabs or post-infection homogenized reproductive tissues. GBS CNCTC 10/84 burdens in vaginal swabs (**A**) or tissues 7 days post-infection (dpi) (**B**) after a 4-week diet course. GBS CNCTC 10/84 burdens in vaginal swabs (**C**) or tissues 7 dpi (**D**) after an 8-week diet course. GBS CNCTC 10/84 burdens in vaginal swabs (**E**) or tissues 2 dpi (**F**) after a 12-week diet course. GBS A909 burdens in vaginal swabs (**G**) or tissues 7 dpi (**H**) after a 12-week diet course. *n*=13-14 (A-B), *n*=8-12(C-D), *n*=10(F-F), *n*=6-7(G-H). Data represent 2 independent experiments (A-H). Points indicate individual samples, and lines or curves indicate medians. Data were analyzed by two-way ANOVA with Benjamini, Krieger and Yekutieli correction and false discovery rate (FDR) set at 5% (A-H) and no significant differences were detected. Supplemental to Figure 1.

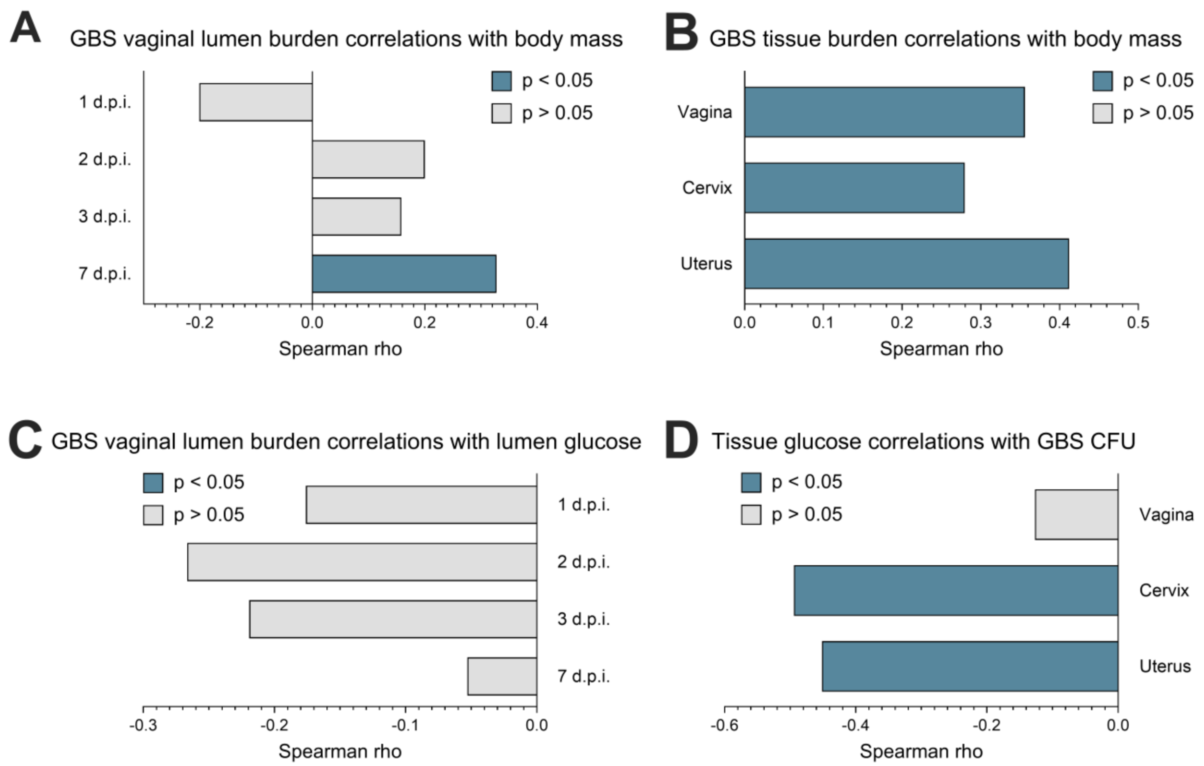


**Figure S2: Body mass and reproductive tract glucose correlations with GBS burdens**. Spearman correlations of body mass with GBS vaginal lumen CFU (**A**) or GBS tissue burdens at 7 days post-inoculation (dpi) (**B**). **C**) Spearman correlations of baseline (12 week) vaginal lumen glucose with GBS vaginal lumen CFU post-inoculation. **D**) Spearman correlations of 7dpi tissue glucose levels with their respective GBS CFU 7dpi. *n*=16-28. Data represent 4 (A-B, D) and 3 (C) independent experiments. Data were analyzed by Spearman correlation with Benjamini-Hochberg FDR correction. Supplemental to Figure 1.

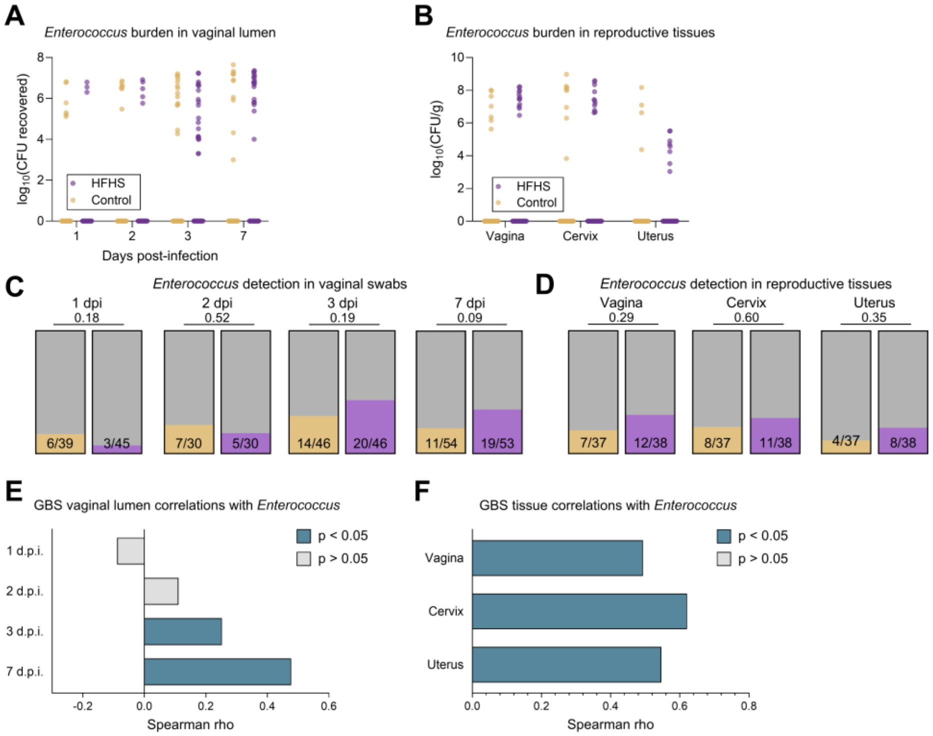


**Figure S3: *Enterococcus* in vaginal lumen and reproductive tract tissues**. **A**) *Enterococcus* burdens from post-infection vaginal swabs. **B**) *Enterococcus* burdens from post-infection homogenized reproductive tissues at 7 days post-infection (dpi). **C**) Proportions of mice in each group (control: yellow, HFHS: purple) with *Enterococcus* detected in vaginal swab samples. **D**) Proportions of mice in each group with *Enterococcus* detected in reproductive tract tissues at 7 dpi. Spearman correlations between GBS CFU and *Enterococcus* CFU in vaginal swabs at all timepoints (**E**) and in reproductive tissues at 7 dpi (**F**). *n*=30-54 (A,C,E), *n*=37-38 (B,D,F). Data represent 4 independent experiments. Points indicate individual samples, and lines indicate medians. Data were analyzed by two-way ANOVA with Benjamini, Krieger and Yekutieli correction and false discovery rate (FDR) set at 5% (A-B) or Fisher's Exact test (C-D) and no significant differences were detected. Data were analyzed by Spearman correlation with Benjamini-Hochberg FDR correction (E-F). Supplemental to Figure 2.

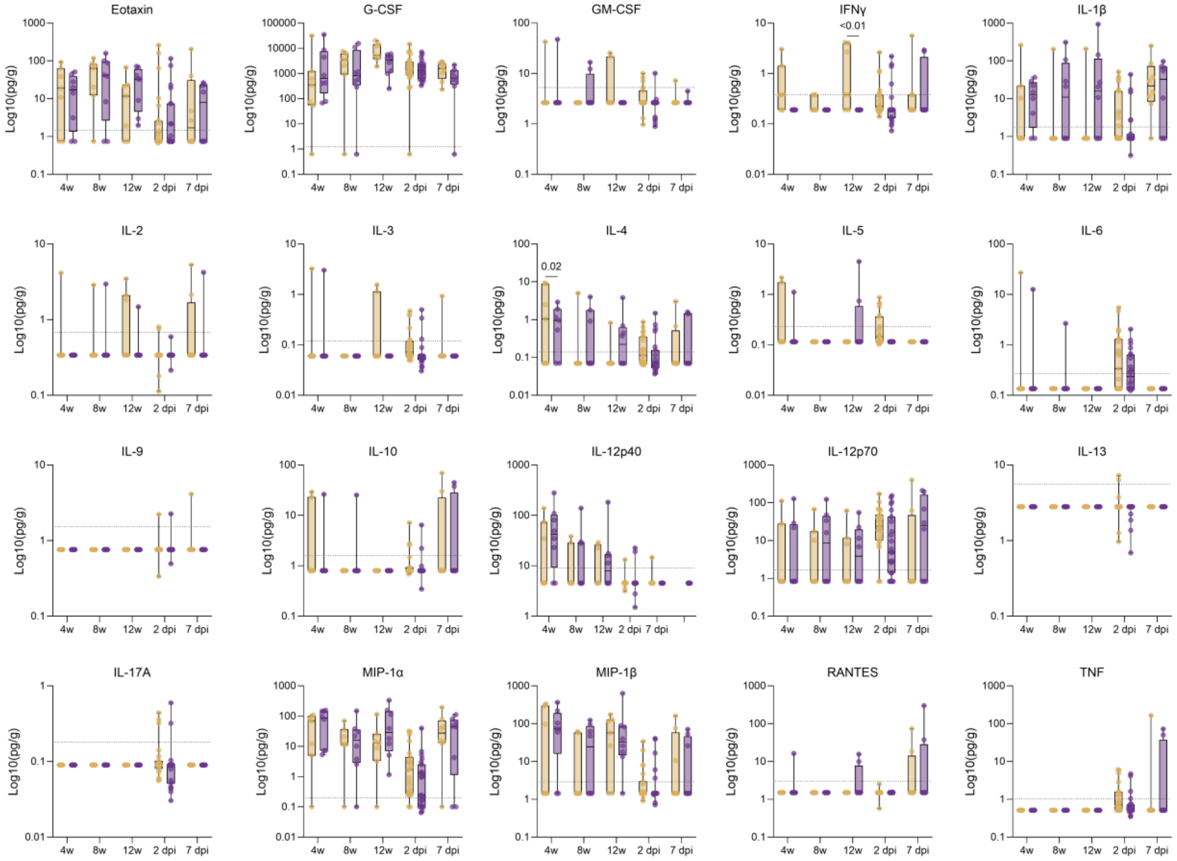


**Figure S4: Vaginal cytokine concentrations at all timepoints**. Cytokine concentrations of vaginal lavage fluid taken pre- and post-infection, normalized to total protein. Data represent 1 (pre-infection and 7 dpi timepoints) or 3 (2 dpi timepoint) independent experiments. Horizontal dotted lines represent lower limit of detection (LLOD). Concentrations that were below the LLOD were extrapolated from standard curves where possible. Where extrapolation was not possible, the values were imputed as half the LLOD. Supplemental to Figure 3.

**Supplemental Table 1. Supplemental data for 16S rRNA gene amplicon sequencing.** ASV counts with their associated taxonomy and consensus sequence, ASVs removed during decontamination, and results of differential abundance and presence analyses. Data represent 2 independent experiments, *n*=8-16 per timepoint. Supplemental to Figure 2.

**Supplemental Table 2. Source data and Spearman correlations for metabolic markers, GBS burdens, and cytokines at various timepoints.** Spearman correlations between metabolic markers and cytokines at indicated timepoints, Spearman correlations between GBS CFU and cytokines at indicated timepoints, and source data for GBS CFU and cytokine (pg/g of total protein) assays. Supplemental to Figures 3, 4, and S4.
